# Electrophysiology in nanoscale compartments

**DOI:** 10.1101/2025.08.29.673146

**Authors:** Madeleine R. Howell, Rosalind J. Xu, E. Cohen Adam

## Abstract

Voltage-gated ion channels play important roles in many membrane-enclosed structures, including synaptic vesicles, endosomes, mitochondria, chloroplasts, viruses and bacteria. Here we study how compartment size and channel gating interact to shape voltage dynamics and ion content in sub-micron structures. In small compartments, assumptions underlying conductance-based (Hodgkin-Huxley type) models of membrane voltage must be relaxed: [1] stochastic gating of individual ion channels can quickly and substantially change membrane voltage; [2] these changes can equilibrate faster than channel state transitions; and [3] ionic currents, even through as few as two channels, can substantially alter ionic concentrations. We adapted conductance-based models to incorporate these effects, and we then simulated voltage dynamics of small vesicles as a function of vesicle radius and channel density. We identified regimes in this parameter space with qualitatively distinct dynamics. We then performed stochastic simulations to explore the role of Na_V_1.5 in maturation of macrophage endosomes. The stochastic model predicted dramatically different dynamics compared to a deterministic approach. Electrophysiology of nanoscale structures can be very different from larger structures, even when ion channel composition and density are preserved.

**SIGNIFICANCE:** With tools of optical electrophysiology, one can measure and perturb membrane voltage in sub-micron structures. Recent experiments in organelles, dendritic spines, and bacteria motivate a re-examination of basic assumptions about bioelectrical phenomena in these compartments. This paper provides a framework for predicting and interpreting bioelectrical dynamics in small structures.

## INTRODUCTION

Ion channels and pumps operate not only in large, patch-clamp-accessible cells, but also in many small membrane-enclosed structures, including synaptic vesicles (1, 2), endosomes (3), lysosomes (4), mitochondria (5), chloroplasts (6), and diverse microbes and viruses (7–11) (**Fig. 1A**). Electrochemical gradients in these compartments are integral to cell physiology: they drive ATP synthesis in mitochondria (12); regulate acidification and cargo processing in secretory and endocytic organelles (2, 13, 14); and contribute to pH- and voltage-dependent stages of viral entry (15, 16). In bacteria, voltage- and mechano-sensitive ion channels participate in homeostasis of membrane potential, osmotic pressure, and ionic balance (8, 17). Yet, despite their ubiquity and functional importance, the electrophysiology of sub-micron compartments has historically been difficult to interrogate because of their small size, intracellular location, and (in microbes) the presence of a cell wall (18).

**FIGURE 1.**
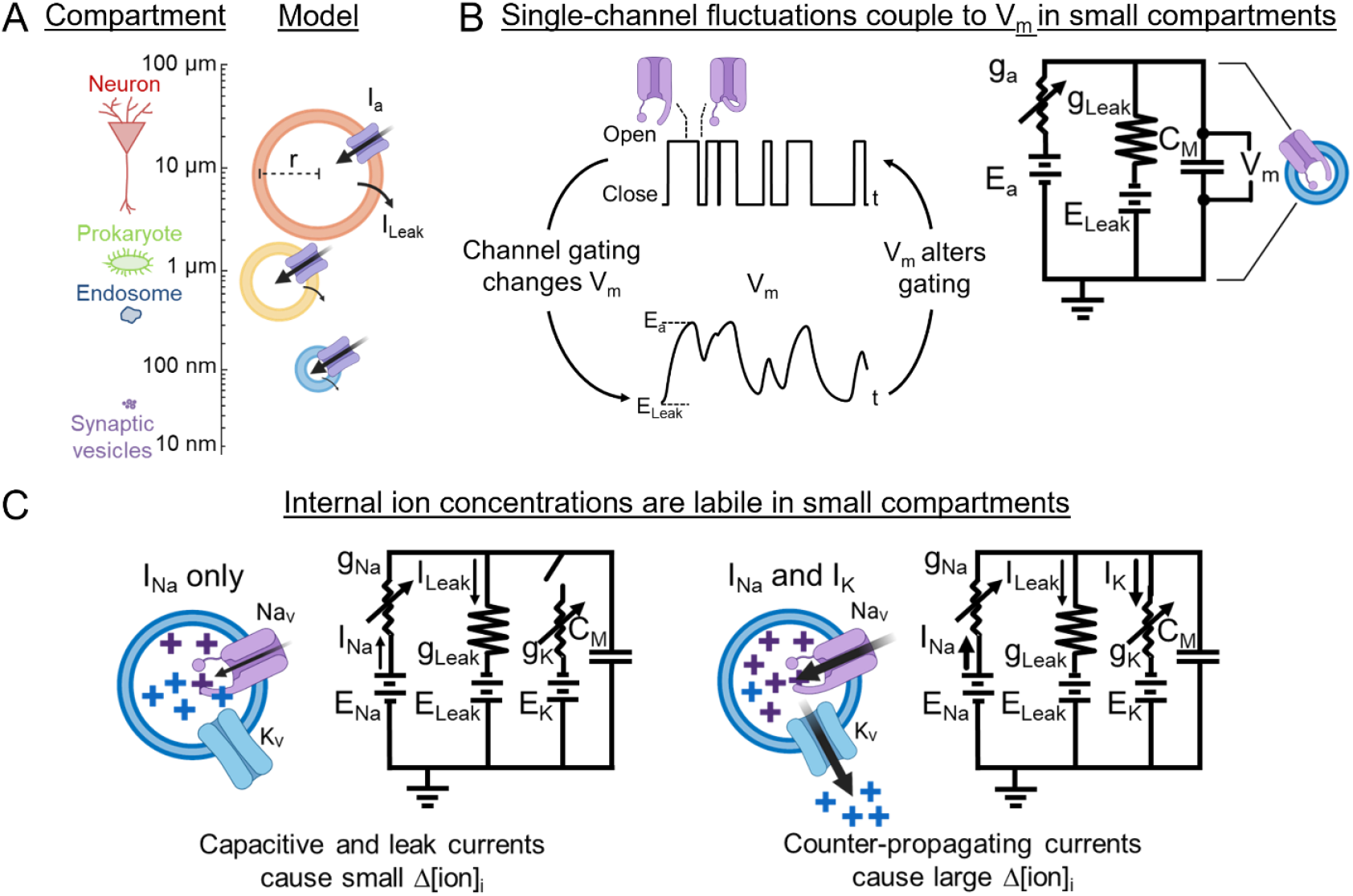
Small compartments have distinctive electrophysiology. (**A**) Bioelectrical dynamics occur over a wide range of spatial scales. Excitable compartments (left) and model vesicles with simplified geometry (right). (**B**) In a small compartment, stochastic gating of a single channel changes *V*_m_, which then affects channel gating. Changes in *V*_m_ alter the duration of individual channel open events and introduce a memory into the dynamics that leads to non-Markovian dynamics. In the circuit diagram, the size of the leak resistor indicates *g*_Leak_ ≪ *g*_a_. (**C**) Coincident activation of channels with different reversal potentials can drive spontaneous depletion of ionic concentration gradients. Upon Na_V_ opening, *I*_Na_ charges the membrane capacitance and then maintains steady-state flow out via *I*_Leak_ (left panel). For realistic parameters, the capacitive charging current has little effect on internal Na^+^ concentration. When Na_V_ and K_V_ opening coincide, inward *I*_Na_ is balanced by outward *I*_K_ until *E*_Na_ and *E*_K_ are equal (right panel). This current drives large changes to internal ion concentrations. Abbreviations: *g*_a_: single-channel conductance of an arbitrary channel; *g*_Na_: Na_V_ single-channel conductance; *g*_Leak_: leak conductance; *g*_K_: K_V_ single-channel conductance; *E*_a_: reversal potential of an arbitrary channel; *E*_Na_: Na^+^ reversal potential; *E*_Leak_: leak reversal potential; *E*_K_: K^+^ reversal potential; *I*_Na_: Na_V_ current; *I*_Leak_: leak current; *I*_K_: K_V_ current; *C*_M_: membrane capacitance; *V*_m_: membrane potential.

This experimental barrier is now largely breached. Genetically encoded voltage indicators (18– 23) and small-molecule dyes (24–27) enable *in situ* voltage measurements in small and/or intracellular membranes (28, 29), with targeted implementations in mitochondria (30, 31), dendritic spines (32), distal astrocytic processes (33), lysosomes (14, 34), and bacteria (35, 36). Complementing protein and dye indicators, DNA-based probes have directly reported organelle electrical and ionic state variables in live cells: an organelle-targeted “voltmeter” reported absolute membrane potentials across multiple organelle classes (34); single-organelle Na^+^ mapping revealed pronounced heterogeneity and maturation-dependent shifts in lumenal sodium (37); and an organelle-localized reporter indicated that voltage-gated K^+^ channels canonically associated with the plasma membrane can be functionally active on trafficking organelles, with measurable consequences for lumenal K^+^ (38).

Together, these experiments raise mechanistic questions that are difficult to answer with classical electrophysiology models: how are organelle voltages and Na^+^/ K^+^ gradients established and maintained when copy numbers of channels/transporters are small (39–41)? When organelle-to-organelle variability is large (37), does this reflect deterministic maturation-dependent shifts, stochastic variations in channel number, or rare but consequential stochastic gating events? And how do transient channel openings during trafficking reshape lumenal ion composition, potentially modulating downstream processes such as acidification, cargo sorting, and maturation (13, 14)?

Answering these questions presents a challenge for the standard theoretical framework for membrane voltage—the Hodgkin-Huxley (HH) family of conductance-based models—which rests on approximations appropriate for large membranes. First, HH-type descriptions are mean-field: conductances represent ensemble averages over many channels, and fluctuations are commonly treated as small perturbations around the mean. In the regime of finite-but-large channel number *N*, channel noise is often well approximated by a Gaussian perturbation to conductance (or gating variables), yielding Langevin/Fokker-Planck descriptions in which voltage performs a diffusion-like biased random walk about a stable fixed point and spiking can be described as a Kramers escape process (42–49). A substantial literature has explored how membrane area and morphology filter or amplify this conductance noise and how it impacts spike timing and reliability (43, 50– 52). These approaches have been powerful precisely because individual channel events contribute only weakly to *V*_m_, and because averaging over many channels smooths the dynamics.

Sub-micron compartments can instead inhabit a qualitatively different regime in which voltage dynamics are dominated by discrete single-channel gating events, not by Gaussian fluctuations around a mean conductance. Two basic scalings drive this transition. (i) Membrane leak conductance scales with membrane area, so the same single-channel current produces a larger steady-state Δ*V*_m_ in a smaller compartment; in sufficiently small structures, opening of a single channel can drive *V*_m_ close to that channel’s reversal potential. (ii) The membrane charging time in small structures can become fast compared to the characteristic channel state-transition times, so channels “feel” their own impact on *V*_m_ during an opening event. This self-action breaks the separation between channel kinetics and voltage dynamics that underlies diffusion approximations: instead of nearly continuous, Gaussian-like voltage noise, the system exhibits jump-like trajectories with history-dependent effective kinetics (**Fig. 1B**). In this regime, it is no longer sufficient to treat stochasticity as a small, additive perturbation to HH dynamics.

A second HH assumption also can break down in small compartments: that (except for Ca^2+^) ionic concentrations are effectively constant on the timescale of electrical activity. Extensions of conductance-based models to include dynamic ion concentrations and pumps have been developed for neurons and cardiac tissue (53, 54), but typically in parameter regimes where concentration changes are slow compared to individual spikes. In contrast, the same small volumes that amplify voltage responses also amplify concentration changes driven by ionic fluxes; moreover, coincident opening of channels with distinct reversal potentials can produce large counter-propagating ionic currents even when net membrane current is small, rapidly dissipating gradients in ways not captured by fixed-reversal-potential models (**Fig. 1C**). The relevance of this issue is underscored by emerging measurements showing that organellar ionic contents can be variable and responsive to channel activity (37, 38). Thus, a theory aimed at interpreting voltage and ion measurements in small compartments must simultaneously confront (a) discrete stochastic gating that directly controls *V*_m_, (b) feedback from *V*_m_ onto channel gating on the timescale of single gating events, and (c) time-dependent driving forces arising from labile ionic gradients.

Here, we adapt conductance-based electrophysiology to explicitly incorporate these small-compartment effects. We treat channel gating as discrete stochastic jump processes (Markov state models) coupled self-consistently to membrane voltage, and we update ionic concentrations (and hence reversal potentials) as currents flow in nanoscale volumes. We use this framework to map regimes of qualitatively distinct dynamics as a function of vesicle size and channel density, to identify when deterministic HH-like descriptions approximate stochastic dynamics and when they fail, and to predict new behaviors—such as voltage “memory” and gradient-depletion–mediated refractoriness—that emerge when single-channel events dominate. Finally, motivated by experimental reports of Na_V_1.5 in macrophage endosomes and its proposed role in acidification (3), we apply the stochastic, concentration-aware framework to endosome maturation, illustrating how rare channel openings can produce outsized physiological consequences in a small compartment. While we primarily analyze the roles of Na_V_ and K_V_ channels, the methodology is not limited to the canonical HH channels, but can apply to the many organelle-specific channels and transporters (55). Together, these results provide a quantitative framework for emerging measurements of voltage and ionic composition in organelles, microbes, and other small compartments, and they suggest experimentally testable signatures of the single-channel-dominated regime.

## METHODS

All simulations were performed in MATLAB.

### Single-channel vesicle model

We examine the interaction between stochastic channel gating and membrane voltage (*V*_m_) using a single 14 pS conductance voltage-gated Na^+^ channel (Na_V_) in the membrane of a spherical vesicle. The vesicle also contains a non-specific and deterministic leak conductance proportional to vesicle area (*A*_m_) with 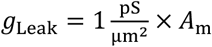 and the leak reversal potentials (*E*_Leak_) given below. Simulations of vesicle *V*_m_ are initialized at *V*_m_ = *E*_Leak_. Because unidirectional single-channel currents do not substantially alter internal electrolyte concentrations, Na^+^ and leak reversal potentials are not updated in simulations of single channel stochastic Na_V_ trajectories. The Markovian model of stochastic Na_V_ gating is described below (**Markovian dynamics for voltage-gated ion channels)** and in **Note S1**. To examine the role of window currents on channel behavior, we modify Na_V_ kinetics to yield a model with substantial window current 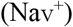 as well as negligible window current 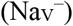. These modifications are described in **Note S1**.

We simulate [1] *V*_m_ in response to a single channel opening (**Fig. 2A**); *E*_Leak_ = −93 mV [2] time-to-close of a channel initialized in the open state (**Fig. 2C**); *E*_Leak_ = −25 mV, and [3] spontaneous trajectories of channel gating and *V*_m_ using either the default Na_V_ model (**Fig. 2B**), *E*_Leak_ = −25 mV; or the modified window current models (**Fig. 2G**), *E*_Leak_ = −60 mV. *E*_Leak_ of −25 mV in [2] and [3] was chosen such that *V*_m_ does not transit the window current regime, to isolate the effect of depolarization on channel kinetics and memory. *E*_Leak_ of −60 mV in [3] was chosen such that channel activation forced *V*_m_ to transit the window regime.

**FIGURE 2.**
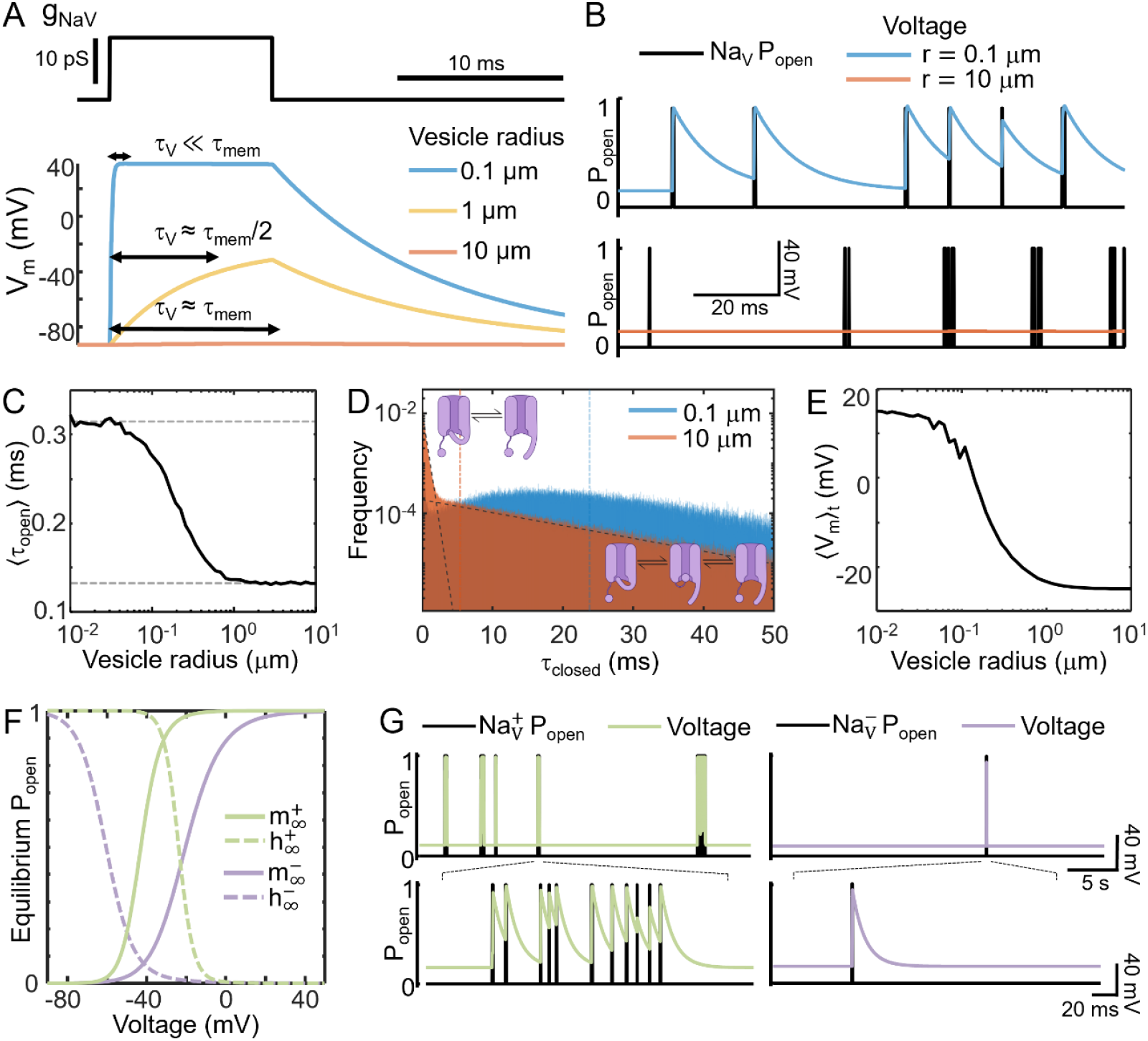
Reciprocal interactions between single-channel gating and *V*_m_ fluctuations in small compartments. (**A**) Simulated *V*_m_ in response to a single Na_V_ channel opening in vesicles of varying size. *τ*_V_ is the time constant of the voltage response when the channel is open; *τ*_mem_ is the intrinsic membrane time constant (when the channel is closed). (**B**) Spontaneous Na_V_ gating (black) and *V*_m_ for *r* = 10 μm (orange) and *r* = 0.1 μm (blue). Leak reversal potential, *E*_Leak_ = −25 mV. (**C**) Mean Na_V_ open-state dwell time as a function of vesicle radius; average of n = 10^4^ simulations of a Na_V_ channel initialized in the open state with *V*_m_(*t* = 0) = *E*_Leak_. Dashed lines show open-state dwell time when *V*_m_ is clamped at *V*_m_ = *E*_Leak_ (bottom) and *V*_m_ = *E*_Na_ (top). (**D**) Closed-state dwell time histograms for voltage trajectories in vesicles with *r* = 10 μm and 0.1 μm, calculated as in **B**; simulations 5,000 s. Black dashed lines indicate linear fits to the fast 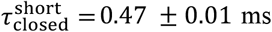 (95% confidence) and slow 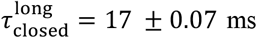 (95% confidence) closed state dwell times for the *r* = 10 μm vesicle. Orange and blue vertical lines indicate mean *τ*_closed_ for *r* = 10 μm (5.4 ± 12 ms) and 0.1 μm (24 ± 19 ms) vesicles (mean ± std). (**E**) Time-average membrane voltage during spontaneous single-channel gating trajectories as in **B**. Each data point is an average of five 50 s-long simulations. (**F**) Steady-state activation (solid trace) and inactivation (dashed) curves for a Na_V_ model with a large window current ( 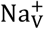, light green) and a Na_V_ model with a small window current ( 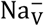, light purple). (**G**) Spontaneous Na_V_ gating (black) and *V*_m_ for the 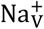 model (left, light green) and the 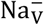 model (right, light purple) in **F**, for *r* = 0.1 μm vesicles at *E*_Leak_ = −60 mV. The 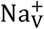 model voltage chatters between epochs of high-frequency oscillation and quiescent epochs.

Annotated code for simulating the dynamics in **Fig. 2B, C** and **G** is available at: https://github.com/adamcohenlab/howell2025small-scale-ephys/tree/main/code-single_NaV_vesicles

### Hodgkin-Huxley (HH) -type vesicle model

We examine the ensemble effects of stochastic and counter-propagating currents in vesicles with HH-like Na_V_, voltage-gated K^+^ channel (K_V_) and leak conductances (1). While the original HH model does not explicitly consider intracellular ion concentrations, it can be extended to an ion-specific form (53, 56–62). Following the conventions of Hübel *et al*., the ion concentrations and reversal potentials are updated at each time step, a Na^+^/K^+^-ATPase model is added to restore the Na^+^/K^+^ balance, and the non-specific leak current is rendered ion-specific (53, 57). Na_V_ and K_V_ channels are modeled as discrete and stochastic with Markovian dynamics (described in **Markovian dynamics for voltage-gated ion channels**). All other conductances are modeled as continuous and deterministic. We assume similar mobilities for Na^+^ and K^+^ ions, so the contribution of each ion to the injected current is proportional to its mole fraction in the extravesicular solution. The formulation is:

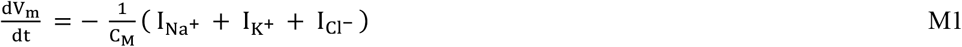

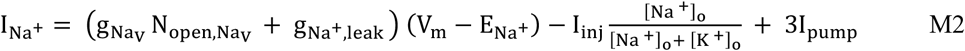

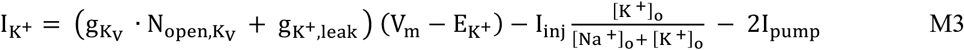

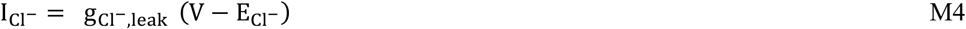

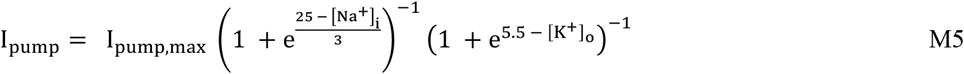

Here *V*_m_ is the membrane potential, *C*_M_ is the membrane capacitance, *E*_ion_ is the reversal potential; *I*_ion_ is the ionic current, *I*_inj_ is the injection current, *I*_pump_ is the Na^+^/K^+^-ATPase current; 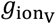 is the conductance of a single voltage-gated channel, 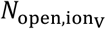 is the number of open channels, *g*_ion,leak_ is the leakage conductance, [ion]_i_ is the internal ion concentration, and [ion]_o_ is the external concentration. All vesicles are modeled as spheres. The electrophysiological parameters and initial conditions of the HH-like vesicle model are in **Table S1** and **Note S1**.

Annotated code to simulate the dynamics in **Fig. 3** and **4** is available at: https://github.com/adamcohenlab/howell2025small-scale-ephys/tree/main/code-HH_neuron_vesicles

**FIGURE 3.**
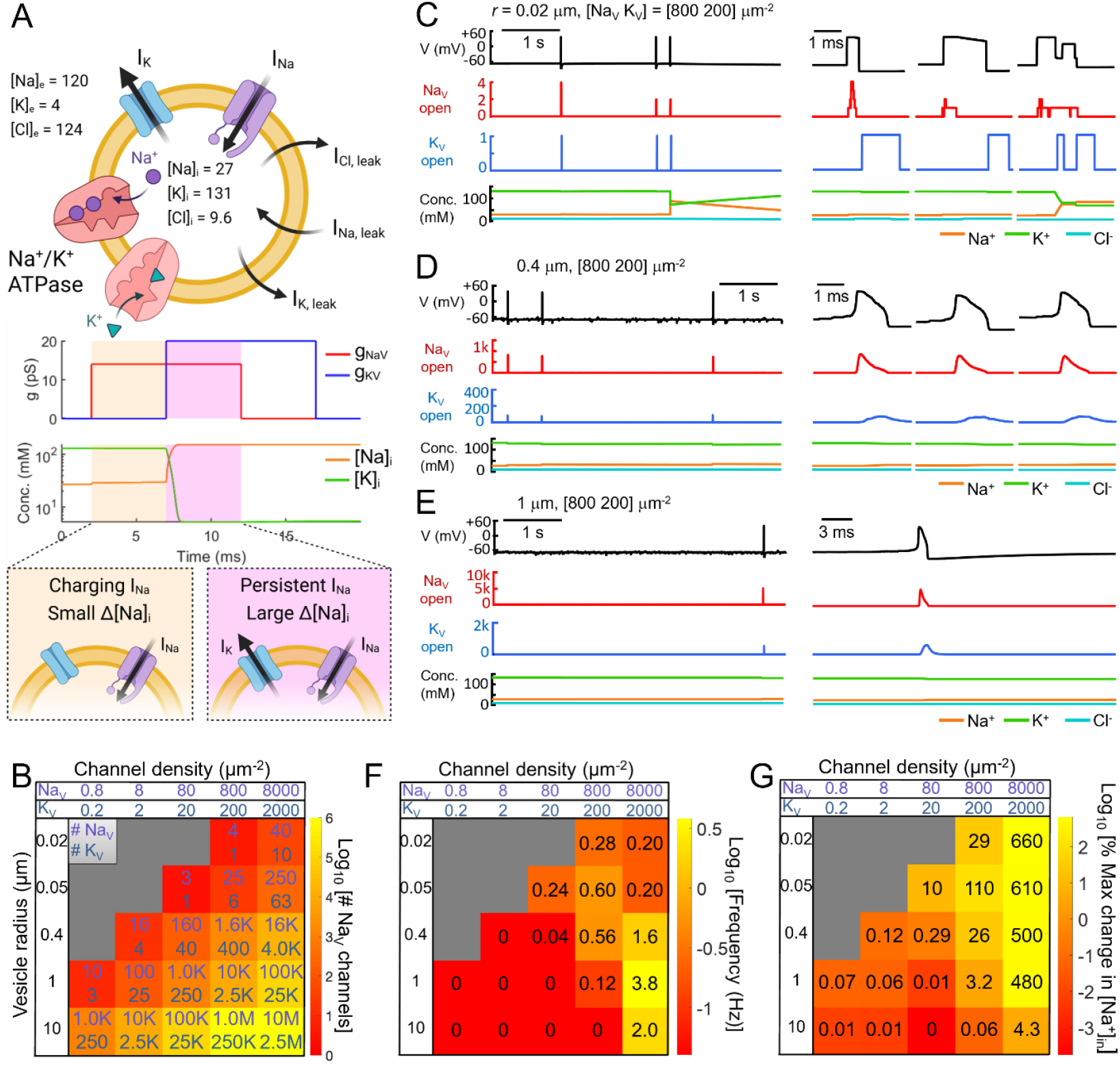
Spontaneous voltage dynamics in small compartments. Voltage and ion concentration dynamics were simulated in vesicles with different sizes and channel densities. (**A**) Top: Model vesicle with HH-type conductances and Na^+^/K^+^-ATPase. All concentrations in mM. Beige panels: Activation of an individual Na_V_ channel does not appreciably change internal [Na^+^]. Pink panels: Concurrent Na_V_ and K_V_ activation depletes [Na^+^] and [K^+^] concentration gradients until the Na^+^ and K^+^ reversal potentials become equal. Plots are for an *r* = 0.02 μm vesicle. (**B**) Range of vesicle sizes and channel densities investigated. Gray regions correspond to size and density combinations with less than one K_V_ channel per vesicle. (**C**) Left: 5 s simulation of a vesicle with *r* = 0.02 μm and [Na_V_, K_V_] = [800, 200] μm^2^, corresponding to 4 Na_V_ channels and 1 K_V_ channel. Right: close-up of the three spontaneous events. In the third event, Na_V_ and K_V_ channel activation overlapped in time, leading to partial dissipation of the ionic gradients. (**D**) Left: 5 s simulation of a vesicle with *r* = 0.4 μm and [Na_V_, K_V_] = [800 200] μm^2^, corresponding to 1,600 Na_V_ channels and 400 K_V_ channels. Overlapping spontaneous Na_V_ activations were sufficient to trigger well-defined spikes. Changes in internal ionic composition were small, on account of the larger vesicle size compared to **C**. (**E**) 5 s simulation of a vesicle with *r* = 1 μm and [Na_V_, K_V_] = [800 200] μm^2^, corresponding to 10,000 Na_V_ and 2,500 K_V_ channels. Due to the larger vesicle size, the influence of each gating event diminished compared to the vesicle in **D**. Channel noise led to small fluctuations in baseline voltage, which occasionally triggered spontaneous spikes. (**F**) Color map of average spontaneous spiking frequency as a function of vesicle radius and channel density. (**G**) Color map of the average percent amplitude of internal [Na^+^] concentration changes as a function of vesicle radius and channel density. These fluctuations were driven by overlapping Na_V_ and K_V_ currents. Data in **F** and **G** are the mean of five, 5 s simulations, excluding the *r* = 10 μm and [Na_V_, K_V_] = [8000 2000] μm^2^ vesicle, which was simulated for 500 ms due to computational constraints.

To compare dynamics between deterministic and stochastic simulations, we used the autocorrelation of the spike train. We chose this measure because it was not sensitive to the precise spike timing. Our measure of similarity plotted in **Fig. 4D** was: 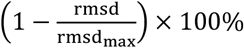, where rmsd is the root-mean-square deviation between the autocorrelation traces, and rmsd_max_ is the maximum rmsd seen across all vesicle sizes and channel densities.

**FIGURE 4.**
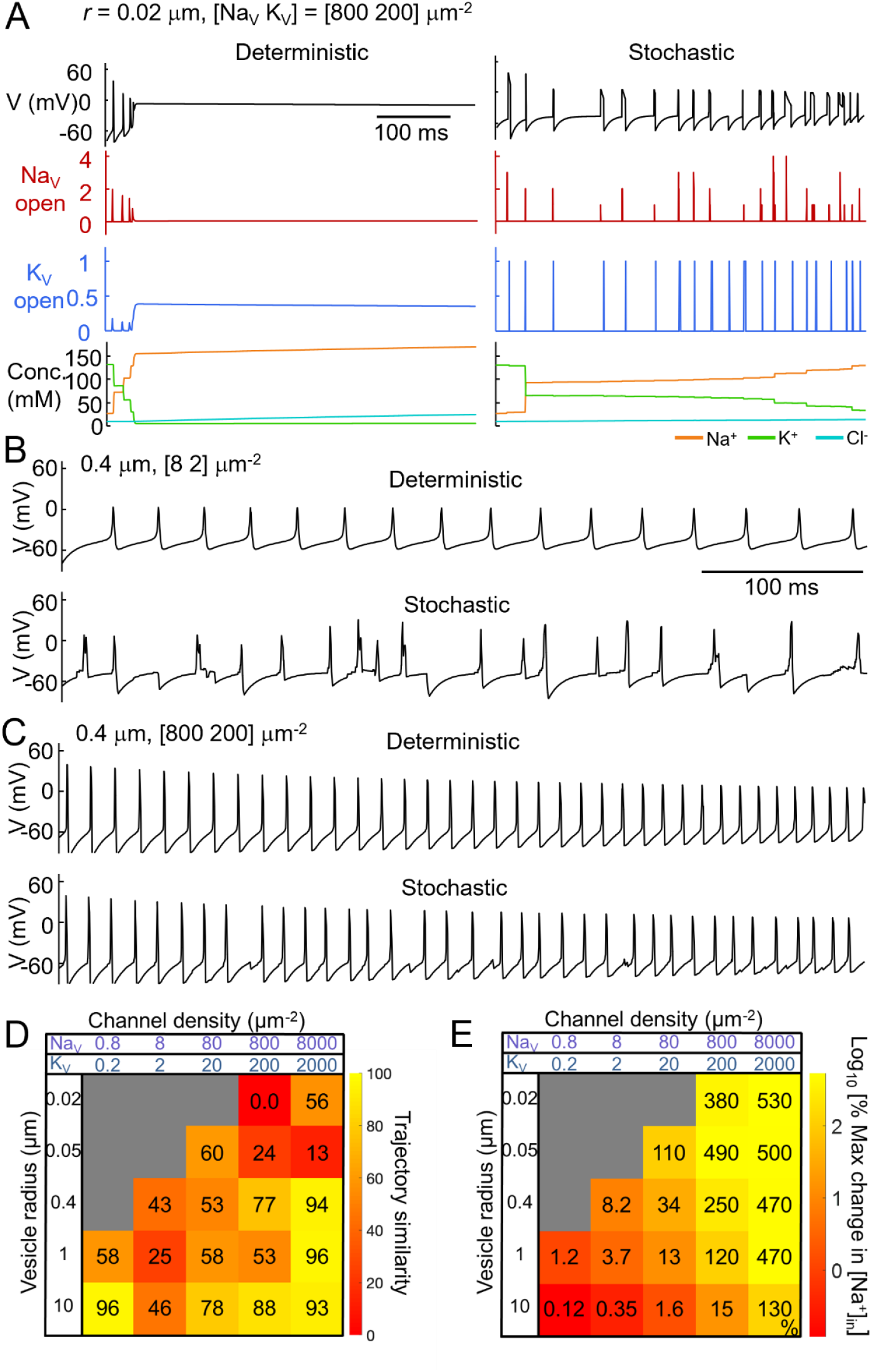
Driven voltage dynamics in small compartments. Simulations were performed as in **Fig. 3**, with the addition of a constant inward current density (Methods). (**A**) At the smallest vesicle size, the deterministic simulation (left) quickly entered depolarization block. In the stochastic simulations (right), fluctuations in channel gating restored the voltage from depolarization block, leading to sustained, but erratic, spiking. (**B**) In an *r* = 0.4 μm vesicle with low channel density, the evoked spikes were variable in waveform due to stochastic variations in the number and timing of channel opening. (**C**) In an *r* = 0.4 μm vesicle with higher channel density, stochastic channel fluctuations caused spike timing to be more variable than in deterministic simulations. (**D**) Color map of the similarity between stochastic and deterministic responses to current injection, as measured by cross-correlation (Methods). In the gray region, there is less than one K_V_ channel. (**E**) Color map of the percent amplitude of internal [Na^+^] concentration changes as a function of vesicle radius and channel density. Data in **D** and **E** are from one, 5 s simulation, excluding the *r* = 10 μm and 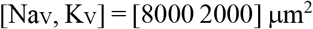 vesicle, which was simulated for 500 ms.

### Endosome model

We study the combined effects of stochastic channel dynamics and ion concentration changes in intracellular endosomes. Carrithers *et al*. reported Na_V_1.5 channels in macrophage late endosomes, and suggested a role in endosome acidification, a part of endosome maturation (3). The minimal endosome model includes Na_V_1.5, vacuolar-type H^+^ ATPase (v-ATPase), and H^+^/Cl^−^ exchange transporter 7 (ClC-7) (63–65). The Na_V_1.5 channels are inserted in an inverted orientation relative to the cell membrane; i.e., the intracellular domain (C-terminus) is facing the outside of the endosome. For ClC-7, a 2:1 Cl^-^/H^+^ coupling ratio is assumed. The formulation is:

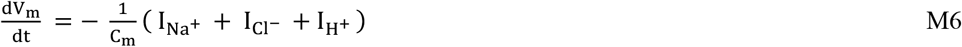

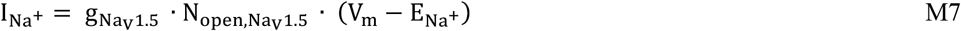

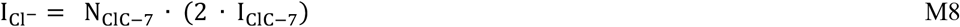

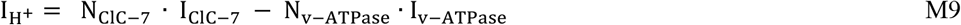

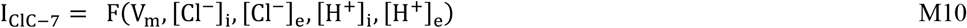

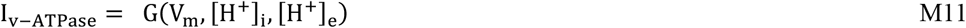

In which I_ClC−7_ is the net current through a single ClC-7 channel, I_V−ATPase_ is the single-transporter v-ATPase current; the formulations for F, G (Eqs. M10 and M11) come from literature (64, 65); N_channel_ is the number of channels. The electrophysiological parameters and initial conditions of the endosome model are in **Table S2**.

Annotated code to simulate the dynamics in **Fig. 5B** and **C** is available at: https://github.com/adamcohenlab/howell2025small-scale-ephys/tree/main/code-endosome

**FIGURE 5.**
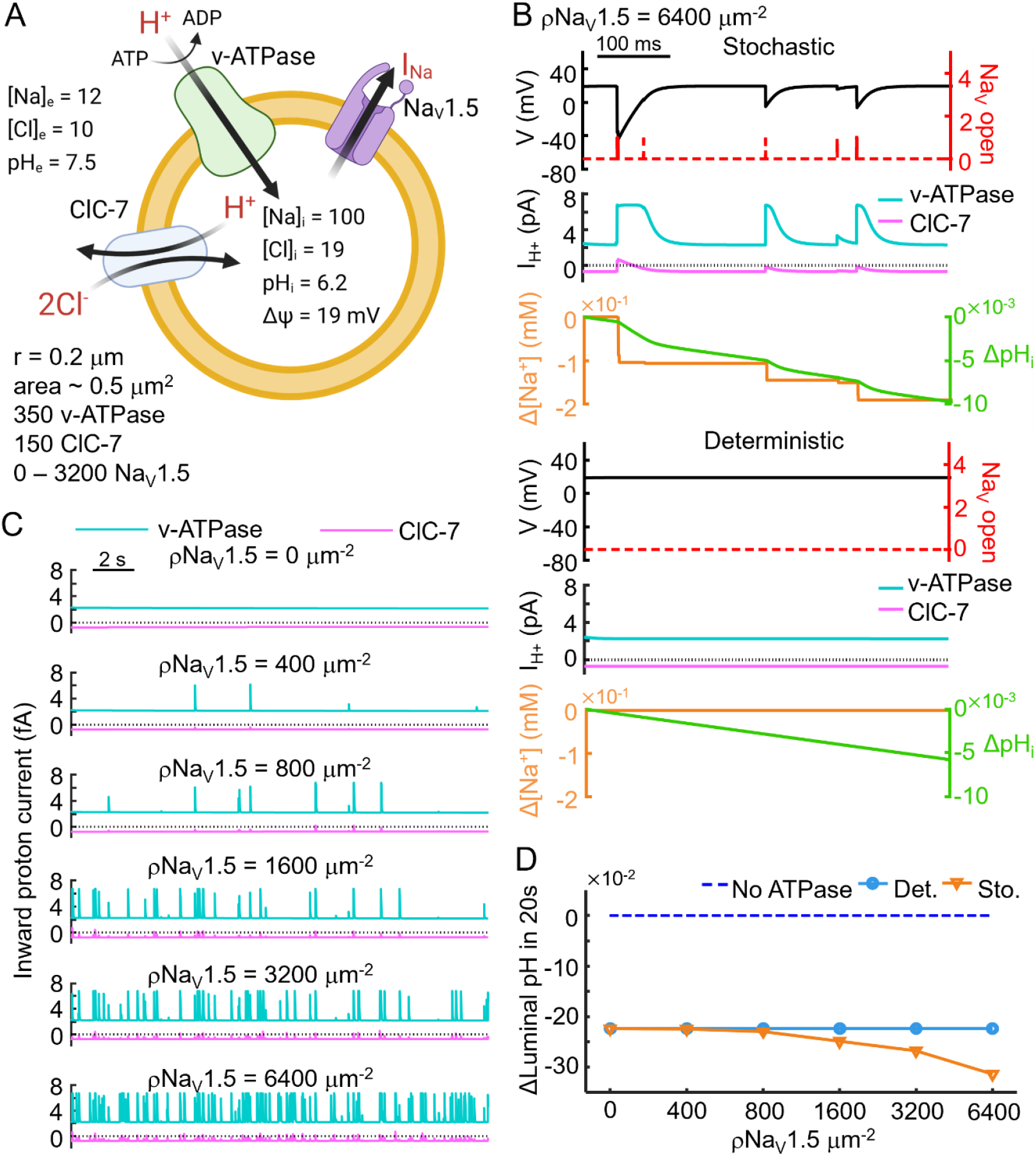
Stochastic simulations of endosome acidification dynamics. (**A**) Model endosome. Concentrations in mM. (**B**) Comparison between the stochastic (top) and deterministic (bottom) dynamics. In the stochastic simulation, spontaneous Na_V_1.5 fluctuations transiently decreased the membrane voltage and increased the acidification rate. (**C**) The effect of Na_V_1.5 activity on endosome acidification increased at higher Na_V_1.5 density. (**D**) Luminal pH in stochastic or deterministic simulations after 20 s, across a range of Na_V_1.5 channel densities. The initial pH is shown as a dashed line.

### Markovian dynamics for voltage-gated ion channels

To facilitate the direct comparison between stochastic and deterministic channel dynamics, Markovian state transition models are used for voltage-gated ion channels in all simulations. The Markovian models for HH-type Na_V_ and K_V_ channels, as well as the Na_V_1.5 channel, come from literature (63, 66–69). The deterministic channel dynamics formulation is:

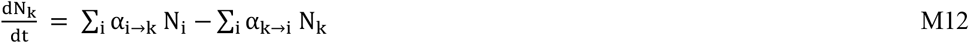

In which N_k_ is the number of channels in state k and α_i→k_ is the transition rate from state i to state k (α_i→i_ = 0). Transition rates for all voltage-gated ion channels are provided in **Note S1** and **S2**.

The Gillespie method is used for the stochastic simulations (67, 69). In short, we define the sum of all the transition rates across all channels in a system as λ:

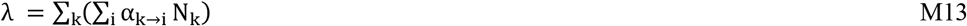

Then the probability that no transition occurs in the system after time *t* follows an exponential distribution:

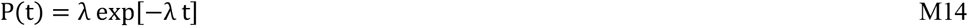

A random number is drawn from this distribution to determine the time elapsed before the next transition. Then, a uniform random number is drawn to determine which type of transition in which type of channel occurs, with weight 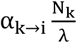 assigned to each type of transition.

### Ion concentration update

The deterministic and stochastic Markovian simulations are performed with intravesicular ion concentrations either fixed or continuously updated. The dynamic ion simulations follow the formalism of Hübel and Dahlem (53, 57). The ionic currents are converted to changes in internal and external concentrations, and the reversal potentials of each ionic species are updated accordingly. For all simulations, the external volume is set to 10^5^ μm^3^; large enough that extravesicular ion concentrations are approximately constant. The ionic composition of the applied current is set proportional to the extravesicular cation ratio. For endosome simulations, a buffering capacity of 40 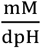 is assumed in updating the intravesicular H^+^ concentration, following literature precedent (70).

### Dynamic choice of simulation step size

To accommodate vesicle sizes and ion channel densities that vary across orders of magnitudes, the simulation step size, *dt*, is chosen dynamically at each step, to satisfy the following criteria: 1) *dt* < 0.001 ms; 2) *dt* < 0.2 τ_V_, where τ_V_ is the characteristic membrane charging time; and 3) In a single step, the internal concentration of each ion does not change by more than 2%.

In driven stochastic simulations, if *dt* chosen from the exponential distribution exceeds the limits set by 1), 2), or 3), then the transition is rejected and the system evolves by the time step dictated by 1), 2), or 3).

In simulations of spontaneous activity in single-channel and HH-type vesicles, constraint 1) was relaxed. This minimized the number of timesteps sampled during quiescent periods in vesicles with few channels. Additionally, in simulations of single-channel vesicles without [ion] update, constraint 3) did not apply and 2) was tightened such that *dt* < 0.01 τ_V_. This enabled accurate calculation of time-averaged *V*_m_.

### Vesicle size and channel density scales

For the HH-type vesicle model, a range of physiologically relevant vesicle sizes and ion channel densities were considered (**Fig. 3B**). The vesicle radii were selected to correspond approximately to a synaptic vesicle (*r* = 0.02 μm) (71, 72); an endosome (*r* = 0.05 μm) (72); a large endosome or vacuole (*r* = 0.4 μm) (73); a prokaryotic cell (*r* = 1 μm); and a eukaryotic cell (*r* = 10 μm). Ion channel densities were selected as follows: [Na_V_ K_V_] = [80 20] μm^-2^ is the approximate density on the squid giant axon, (57, 67) and [Na_V_ K_V_] = [800 200] μm^-2^ is the approximate density on a synaptic vesicle if it contains exactly one copy of the K_V_ channel. On a synaptic vesicle, the copy numbers of each type of ion channel and transporter are often in the single-digit regime (39, 71, 74–76). The ratio of Na_V_ to K_V_ channel was fixed at 4 to 1 across all channel density scales.

For the endosome model, the vesicle radius was 0.2 μm. The density of v-ATPase was estimated to be 700 μm^-2^ and that of ClC-7 was 300 μm^-2^ (64, 70). The density of ClC-7 was adjusted so the initial membrane potential was +19 mV (77). The density of Na_V_1.5 on macrophage late endosomes is not known. We varied this density across a wide range, from 0 to 6400 μm^-2^.

## RESULTS

### Scaling electrophysiology to small systems

#### Single-channel perturbations to membrane voltage

We begin by examining the relationship between membrane voltage and channel dynamics in small systems. **Fig. 1A** depicts the scale of several electrically excitable biological systems. For simplicity, we consider a spherical membrane-bound compartment (a “vesicle”) containing a single ion channel of type *a*, with open-state conductance *g*_a_ (pS) and reversal potential *E*_a_ (mV). Assume the membrane has a specific leak conductance *G*_0_ (pS/μm^2^) and a resting potential *E*_leak_ (mV). Typically *G*_0_ ∼1 pS/μm^2^ for neuronal membranes (53). What is the effect of the single channel opening on the membrane potential, *V*_m_?

If we approximate all the conductances as Ohmic, then a simple circuit analysis shows that (for static ion concentrations, i.e. fixed ion reversal potentials) the change in steady-state membrane potential upon channel *a* opening is:

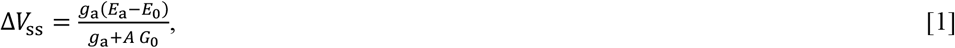

where *A* is the surface area of the vesicle. For example, when *g*_a_ = *A G*_0_, the voltage swing is half its maximum value: Δ*V*_ss_ = (*E*_a_ − *E*_0_)/2. Not surprisingly, the single-channel gating has a substantial effect on the steady-state membrane potential when the unit conductance surpasses the resting leak conductance. **Fig. 2A** illustrates this scenario for a canonical Hodgkin-Huxley Na_V_ channel.

The leak conductance of a vesicle is proportional to its surface area (assuming a high density of leak channels), while the single-channel conductance is independent of the vesicle surface area. Hence, the crossover between charging regimes (i.e. when *g*_a_ = *A G*_0_) corresponds to a characteristic vesicle radius, *r*_c_. For example, for *G*_0_ = 1 pS/μm^2^ and a Na_V_ channel (*g*_a_ = 14 pS) (78), the crossover radius is *r*_c_ = 1.1 μm. For a BK calcium-activated potassium channel (*g*_a_ ∼200 pS) (79), *r*_c_ = 4 μm, while for a homomeric 5-HT_3A_ receptor (*g*_a_ = 1 pS) (80), *r*_c_ = 280 nm.

Upon opening of a single ion channel (**Fig. 2A)**, the RC time constant for the voltage to approach its new steady state is

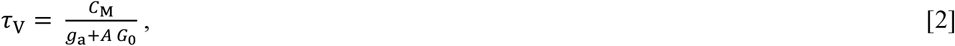

where *C*_M_ = *A c*_0_ is the membrane capacitance (specific capacitance *c*_0_ ∼10 fF/μm^2^), and the denominator is the total membrane conductance. At the crossover radius *r*_c_ defined above, *τ*_V_ = *τ*_mem_/2, where *τ*_mem_ = *c*_0_/*G*_0_ is the intrinsic membrane time constant. For example, for *G*_0_ = 1 pS/μm^2^, *τ*_mem_ = 10 ms and when *r* = *r*_c_, *τ*_V_ = 5 ms. If the single-channel gating events have duration *τ*_open_ ≫ *τ*_V_, then Δ*V*_m_ approaches Δ*V*_ss_ during a single gating event (**Fig. 2A**, blue trace). If *τ*_open_ ≪ *τ*_V_, then the amplitude of the voltage fluctuations is suppressed to Δ*V*_m_ ≈ Δ*V*_ss_*τ*_open_/*τ*_V_ (**Fig. 2A**, yellow trace).

For big vesicles (*r* ≫ *r*_c_ ), single-channel gating changes steady-state voltage by 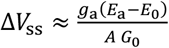 with time constant *τ*_V_ ≈ *τ*_mem_. Since *g*_a_ ≪ *A G*_0_ for big vesicles, Δ*V*_ss_ is a small perturbation to the resting potential (**Fig. 2A**, orange trace). This corresponds to the situation previously analyzed in models of channel stochasticity in neural firing (45–49).

For small vesicles (*r* ≪ *r*_c_), single-channel gating brings the voltage close to the reversal potential of channel *a*, i.e. Δ*V*_ss_ ≈ (*E*_a_ − *E*_0_). In this case the time constant is 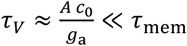. For example, upon activation of a single 14 pS channel in a vesicle of *r* = 0.1 μm, the voltage responds with a time constant *τ*_V_ ≈ 10^−4^ s (**Fig. 2A**, blue trace). When the channel closes, the voltage relaxes back toward the resting potential with time constant *τ*_mem_.

Thus, for sufficiently small structures, the single-channel influence on the membrane voltage can be substantial and the membrane voltage equilibrates faster than the characteristic channel gating times (**Fig. 2B)**. Single ion channels then ‘see’ their own influence on *V*_m_ during a stochastic gating event. This self-action changes the statistics of channel fluctuations.

For example, consider a vesicle containing a single Na_V_ channel (Na_V_ model in **Fig. S1**) and a constitutively open ‘leak’ conductance. Here we assume internal ion concentrations are fixed (an assumption relaxed later) and we initialize the channel in the open state at *V*_m_ = *E*_Leak_. In a large vesicle, Na_V_ opening (gating variables *m*^*3*^: 0 → 1; *h*: 1) is quickly reversed through deactivation (*m*^*3*^: 1 → 0). In a small vesicle, Na_V_ opening depolarizes the vesicle to a voltage which favors the open state. The channel then remains open until open-state inactivation sets in (*h*: 1 → 0). Closing of the channel leads to an increase in the membrane time-constant from *τ*_V_ to *τ*_mem_, so the voltage recovery is slower than the voltage upstroke. **Fig. 2C** shows how voltage self-action prolongs mean Na_V_ open-state lifetimes in small vesicles relative to large vesicles, and **Fig. S2** shows the underlying distributions of Na_V_ open times in small and large vesicles.

In a large vesicle, channel closing after an open-state fluctuation typically resets the gating variables to *m*^*3*^ = 0, *h* = 1. The channel is immediately available to open again. Since single-channel gating has minimal effect on *V*_m_ (**Fig. 2B**, bottom), the channel has constant probability of opening per unit time. Thus, the closed-state waiting-time distribution is exponentially distributed at short times. Occasionally, open-state fluctuations terminate in an inactivated state (*h* = 0). These rare events lead to a long exponential tail to the waiting time distribution (**Fig. 2D**).

In a small vesicle, an open-state fluctuation induces a large depolarization, which continues to act after the channel has inactivated (*h* = 0). The voltage then relaxes back toward *E*_Leak_ with time constant *τ*_mem_ (**Fig. 2B**, top). The time-dependent voltage interacts with the voltage-dependent gating parameters to produce a complex waiting-time distribution (**Fig. 2D**). As a result of the residual depolarization following an open event, the time-average membrane voltage is more depolarized in small vesicles than in large vesicles (**Fig. 2E**).

If the resting potential, *E*_Leak_, is hyperpolarized relative to *E*_Window_, the window current regime is wide enough, and the repolarization is slow enough, then the channel may have a high probability to re-open when the voltage nears *E*_Window_ (**Fig. 2F**, green trace). In trajectories simulated with a Na_V_ model modified to have a large window current, we observed voltage oscillations between *E*_Window_ and *E*_Na_. Occasionally the voltage repolarized below *E*_Window_ without the channel opening, leading to a quiescent epoch with voltage close to *E*_Leak_ (**Fig. 2G**, green trace). A spontaneous opening event would then re-start the oscillatory dynamics. The membrane voltage thus “chattered” between epochs at *E*_Leak_ and oscillations between *E*_Window_ and *E*_Na_ (**Fig. 2F-G**, green trace). This behavior was absent in large vesicles containing the same Na_V_ model. In a Na_V_ model modified to have negligible window current (**Fig. 2F**, purple trace), the voltage repolarized completely to *E*_Leak_ (**Fig. 2G**, purple trace). **Fig. S3** shows the distributions of Na_V_ closed times for these different scenarios. Through large *V*_m_ fluctuations and slow membrane voltage repolarization, small vesicles endow channels with a “memory” of prior gating events, which can lead to a rich repertoire of vesicle size-dependent dynamics.

#### Single-channel perturbations to concentration

Stochastic channel gating can also affect the ionic contents of the vesicle. To change the membrane voltage by an amount Δ*V* requires a flux of charge Δ*Q* = *C*Δ*V*. The ions carrying this charge flow into an internal volume *M*. Thus, the change in concentration, ΔΓ = *C*Δ*V*/(*z F M*), where *z* is the valence of the charge carrier and *F* is the Faraday constant. For a spherical vesicle of radius *r*, we have *C* ∝ *r*^2^ while *M* ∝ *r*^3^, so ΔΓ ∝ 1/*r*, i.e. for a given voltage swing, changes in internal concentration are bigger for smaller vesicles. Numerically, the change in concentration is ΔΓ = 3.1 × 10^−4^ Δ*V*/*z r*, where *V* is measured in mV, *r* in microns, and Γ in mM. For an *r* = 0.1 μm vesicle with a voltage swing Δ*V* = 100 mV, the change in concentration due to charging is ∼0.3 mM/*z*, likely important only for Ca^2+^.

This estimate misses an important factor, however. Simultaneous opening of channels with different reversal potentials leads to steady-state ionic flows. Net current can be approximately zero (so *dV*/*dt* ≈ 0), while distinct ionic species maintain equal and opposite flows, e.g. via balanced Na^+^ influx and K^+^ efflux, leading to substantial changes in concentration.

In **Appendix 1** we analyze this scenario for a persistent Na^+^ current combined with a leak conductance. We assume that voltage equilibrates fast compared to [Na^+^]_in_. This assumption then leads to a rate of change of [Na^+^]_in_ is proportional to the difference in reversal potential between Na^+^ and the leak:

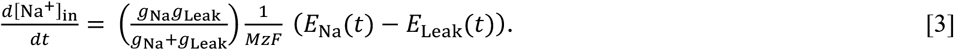

Where *E*_Na_(t) and *E*_Leak_(t) are instantaneous values of the Na^+^ and leak reversal potentials, respectively. The steady-state [Na^+^]_in_ is reached when *E*_Na_(*t*) = *E*_Leak_(*t*). Imposing electroneutrality, i.e. 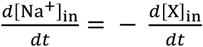, where [X]_in_ is the concentration of the counterion carried by the leak conductance (e.g. Cl^-^ or K^+^), the steady-state value of [Na^+^]_in_ is:

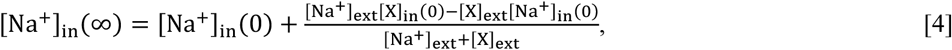

where [X]_in_(0) and [Na^+^]_in_(0) are initial internal concentrations of ions. The external concentrations of ions, [Na^+^]_ext_ and [X]_ext_ are assumed to be fixed. From Eq. 3, we derive the characteristic timescale of changes in [Na^+^]_in_:

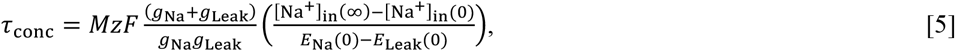

where *E*_Na_(0) and *E*_Leak_(0) are the initial values of the Na^+^ and leak reversal potentials, respectively. Let us consider the case where a single Na_V_ channel ( 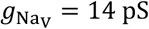, *E*_Na_ = 40 mV) and a single K_V_ channel (e.g. a delayed rectifier K^+^ channel, 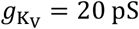, *E*_K_ = -90 mV) are open simultaneously. Then the initial steady-state voltage is -36 mV, the initial steady-state Na^+^ current is *I*_Na_ = -1.1 pA and the initial steady-state K^+^ current is *I*_K_ = +1.1 pA. In a 1 μm diameter vesicle, these currents correspond to 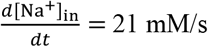. In a 0.1 μm diameter vesicle, 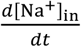 is 000-fold higher, i.e. 21 mM/ms. Thus, a millisecond-duration flicker comprising counter-propagating ionic currents can substantially change the ionic content of a small compartment. **Fig. 3A** shows an example of [Na^+^]_in_ dynamics resulting from a charging current 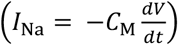 and a persistent current 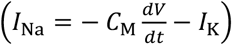 during Na_V_ and K_V_ gating in a small vesicle.

### Electrophysiology in Hodgkin-Huxley-type model vesicles

Next, we explore how ion channel stochasticity, self-action on membrane voltage, and ion concentration dynamics interact to influence electrophysiology in small compartments. We modeled small spherical vesicles with variable densities of Hodgkin-Huxley (HH)-type Na_V_ and K_V_ channels (**Fig. 3A**, Methods). For biologically plausible vesicle sizes and channel densities, channel numbers vary by ∼7 orders of magnitude (**Fig. 3B**). To restore ionic gradients, we added a Na^+^/K^+^-ATPase with a biophysically realistic Na^+^ and K^+^ concentration-dependent pumping rate (57, 81). We added Na^+^, K^+^, and Cl^-^ leak conductances to set a resting potential of -68 mV. The Na^+^/K^+^-ATPase and the leaks were proportional to the membrane area and deterministic. We simulated voltage and concentration dynamics, comparing deterministic and probabilistic Markov models of Na_V_ and K_V_ channel gating (Methods). In all simulations we treated internal ion concentrations as dynamic and external ion concentrations as approximately constant.

#### Spontaneous activity

We simulated trajectories of spontaneous Na_V_ and K_V_ channel gating. As expected, the deterministic simulations produced no spontaneous activity, since both Na_V_ and K_V_ channels remained closed at the resting membrane potential, -68 mV. In stochastic simulations however, Na_V_ and K_V_ channels occasionally fluctuated to the open state. The consequences of these fluctuations depended on vesicle size and channel density.

In small vesicles (*r* < 100 nm), a single Na_V_ channel opening could depolarize the vesicle enough to activate other Na_V_ channels, leading to a spike (**Fig. 3C**). The spike waveforms were highly variable, due to the strong contributions of discrete channel gating events. During some spikes the Na_V_ and K_V_ currents overlapped, driving rapid ion gradient depletion (**Fig. 3C**, inset, right panel). The amount of ion gradient depletion depended on the overlap of the Na_V_ and K_V_ open times. Smaller vesicles supported longer Na_V_ channel openings due to voltage self-action (**Fig 2C**) and higher channel densities created more opportunity for Na_V_ and K_V_ overlap. This gradient depletion caused a long-lasting decrease in vesicle excitability, restored eventually by the activity of the Na^+^/K^+^-ATPase (we assume a constant supply of ATP). This depletion-mediated refractory period is distinct from a typical refractory period associated with ion channel recovery kinetics. Gradient depletion diminished spontaneous spike frequency at high channel densities (**Fig 3F-G**, right column)

In larger vesicles (100 nm < *r* < 1 μm), single-channel events typically induced subthreshold depolarization (Na_V_) and hyperpolarization (K_V_). As channel density increased, these independent single-channel events began to overlap in time, leading to voltage dynamics that resembled an Ornstein-Uhlenbeck process, i.e. diffusion in a parabolic potential. This scenario has previously been studied analytically (67) and numerically (68). When the biased random walk of voltage approached the Na_V_ activation potential, it triggered spikes which resembled classical action potentials (**Fig 3D-E**). The frequency of these events has been modeled as a Kramers escape process (67).

At constant channel density, the spontaneous firing rate showed a non-monotonic dependence on vesicle size (**Fig 3F**). For small vesicles, an increase in size led to an increase in the number of channels. Since a single stochastic channel gating event was sufficient to trigger a spike, more channels led to a higher spontaneous spike rate. For large vesicles, the influence of single-channel gating events on membrane voltage decreased, so spontaneous spikes became less frequent. This non-monotonic trend is similar to results of numerical simulations by Skaugen and Walløe (68). The largest vesicles (*r* > 1 μm) were quiescent at all but the highest channel densities.

#### Evoked activity

We next simulated voltage trajectories in response to constant current injection. To mimic the effect of an electrogenic pump (optogenetic or chemically driven), we made the current proportional to membrane area (*I* ≈ 2.4 × 10^3^ fA/μm^2^). We generated voltage trajectories using both stochastic and deterministic (i.e. HH-like) simulations of channel gating with dynamic internal ion concentrations.

When the number of channels was small, discrete gating events led to substantial deviations between the deterministic and stochastic simulations. Stochastic gating events could lead to spontaneous recovery from depolarization block (**Fig. 4A**), and converted regular spike trains into irregular trains (**Fig. 4B**). As channel number increased, stochastic trajectories came to resemble deterministic trajectories of the same vesicle (**Fig. 4C**). We quantified the similarity between voltage traces for deterministic vs. stochastic simulations via root-mean-square deviation (rmsd) between the autocorrelation functions of the two voltage trajectories (Methods). **Fig. 4D** shows the similarity between deterministic and stochastic simulations as a function of vesicle size and channel density.

We quantified the extent of internal ion concentration changes in the stochastic simulations. The ion gradients were more prone to depletion when the vesicle was smaller and the channel density was higher, as in the simulations in the absence of current injection (**Fig 4E** and **3G**).

### Na_v_1.5 channels and endosome acidification

Finally, we modeled the effects of small-scale electrophysiology on macrophage endosome acidification. This acidification is essential for killing of pathogens engulfed by phagocytosis. Macrophage endosomes express Na_V_1.5, with C-terminus outside the lumen (inverted relative to Na_V_1.5 on the cell membrane). The channel density in endosomes is not known (3), so we simulated vesicles with Na_V_1.5 densities 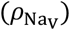 between 0 − 6400 μm^-2^. We assumed radius *r* = 0.2 μm (73) (**Fig. 5A**), initial membrane potential +19 mV (inside relative to outside), initial pH 6.2, and initial internal Na^+^ concentration 100 mM (77, 82). A v-ATPase imports H^+^, acidifying the endosome during maturation (64, 65), and ClC-7 contributes a small outward H^+^ current (64). The electrophysiological parameters and initial conditions are in **Table S2**. The Na_V_1.5 model is described in **Note S2**. The endosomal Na_V_1.5 model accounts for isoform-specific features, including greater steady-state inactivation at moderate to high voltages compared to the canonical HH Na_V_ model.

In deterministic simulations of endosomes with 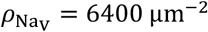, the mean number of open Na_V_1.5 channels was always much less than 1. We rounded the Na_V_ conductance to the nearest integer (in units of the open-state conductance), which rounded to zero. Hence, the Na_V_ channels did not contribute to the deterministic simulations (**Fig. 5B**).

In stochastic simulations at 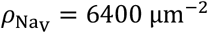, the Na_V_1.5 channels were mostly in the inactive state due to the “felt” voltage of -19 mV (outside relative to inside). Occasionally, a single Na_V_1.5 channel fluctuated to the open state. In the *r* = 0.2 μm vesicle, this induced a decrease in membrane voltage (inside relative to outside) of ∼21±16 mV (mean ± s.d.) since [Na^+^] was higher inside the endosome than in the cytoplasm. The voltage dips recovered with time constant ∼16±2.4 ms (mean ± s.d.). The voltage decrease enhanced v-ATPase activity and suppressed the outward H^+^ current due to ClC-7 activity, speeding up acidification (**Fig. 5B**). The effect of Na_V_1.5 activity on the acidification dynamics became more pronounced as Na_V_1.5 channel density increased, since the fraction of time when there was one Na_V_1.5 channel open was proportional to the number of Na_V_1.5 channels present (**Fig. 5C**).

**Fig. 5D** shows endosomal pH change as a function of the density of Na_V_1.5 channels. In the deterministic simulations, the acidification rate was unaffected by the presence of Na_V_1.5, since the average number of open channels was approximately zero. In the stochastic simulations, however, the acidification rate increased as Na_V_1.5 channel density increased, due to the Na_V_-induced membrane potential fluctuations. Thus, stochastic channel fluctuations are likely relevant to this important physiological process.

## DISCUSSION

This work identifies a qualitative regime change in electrophysiology as membrane-enclosed compartments shrink to sub-micron dimensions. In this regime, (i) single-channel openings can drive large, rapid excursions in membrane potential, (ii) the resulting voltage changes feed back onto channel kinetics during the same opening event, and (iii) ionic currents—especially counter-propagating currents through different channel types—can measurably reshape intralumenal ion concentrations during single gating events. By combining Markov-state channel gating with self-consistent voltage dynamics and ion-concentration updates, we provide a unified framework that connects classical conductance-based descriptions to the single-molecule limit.

A central implication of our work is that small compartments need not behave like miniature versions of excitable cells. In the classical setting (large membrane area and many channels), channel noise is well captured as an approximately Gaussian perturbation around a mean conductance, enabling diffusion/Langevin and Kramers-type descriptions of rare excursions and spontaneous spikes (43, 45–49, 66–68). In contrast, when channel copy numbers are in the single-digit to tens, and leak conductance and capacitance are correspondingly small, voltage trajectories are inherently jump-like: individual gating events can move *V*_m_ a substantial fraction of the way toward a reversal potential, and the ensuing self-action can prolong or curtail dwell times depending on the voltage-dependence of the transition rates. This feedback effectively endows the system with history dependence (“memory”) even when the underlying channel model is Markovian, and it can generate dynamics—such as intermittency or “chattering” in channels with appreciable window currents—that have no direct analog in larger systems.

A second implication concerns ionic homeostasis. In large cells, voltage dynamics are often computed with fixed reversal potentials (except for Ca^2+^), and concentration dynamics enter only as slow modulators (53, 54). In small volumes, this separation can break down. Our simulations highlight a particularly important case: coincident opening of conductances with different reversal potentials can drive large counter-propagating ionic fluxes while producing little net capacitive current. This flux can rapidly dissipate Na^+^/K^+^ gradients, producing a depletion-mediated refractory period that is mechanistically distinct from classic refractoriness set by channel inactivation and recovery. This predicted mechanism suggests a plausible interpretation for recent organelle measurements, where endosome-to-endosome variability in lumenal Na^+^ and maturation-dependent Na^+^ shifts are substantial (37), and where K^+^ channel activity on trafficking organelles measurably changes lumenal K^+^ (38). In particular, our results motivate the hypothesis that some of the observed heterogeneity could arise from rare overlaps of channel openings that transiently reshape ionic gradients.

Our endosome simulations provide a complementary example of how discrete gating can matter even when the mean open probability is extremely small. For Na_V_1.5 in macrophage late endosomes (3), the deterministic (mean-field) description can effectively “round away” the channel contribution when the expected number of open channels is ≪ 1. The stochastic model, however, predicts that rare single-channel openings can generate transient voltage excursions that systematically bias electrogenic transport (v-ATPase, ClC-7), accelerating acidification. This points to a broader principle: in nanoscale compartments, physiological impact may be controlled by event statistics (amplitude, dwell-time distributions, and coincidence rates) rather than by ensemble- or time-averaged conductances.

These considerations suggest several experimentally testable predictions and design principles for organelle and microbial electrophysiology. First, voltage and ion measurements at single-organelle resolution (34, 37, 38) should exhibit non-Gaussian signatures—e.g., heavy-tailed distributions of voltage deflections and state-dependent dwell times—that become more distinct as compartments become smaller or channel copy number decreases. Second, simultaneous measurements of *V*_m_ and lumenal counterions (e.g. [Na^+^] or pH in endosomes) should reveal transient covariance consistent with counter-propagating flux episodes: large voltage events with disproportionately large concentration changes when multiple channel types overlap. Third, perturbations that change organelle size, channel abundance, or kinetics (e.g., channel blockers or mutants that modify window current) should shift systems between qualitatively distinct dynamical regimes in predictable ways, offering a path to infer otherwise inaccessible parameters such as channel copy number per organelle.

Several extensions could broaden the model’s applicability. Many small compartments are not closed: dendritic spines, nodes of Ranvier, and subcellular invaginations exchange ions with a larger reservoir through narrow necks or diffusion-limited pathways (41, 83, 84). Incorporating diffusive coupling to cytosol or extracellular space would allow the same framework to address when local ionic depletion is buffered by exchange versus when it becomes functionally consequential. Similarly, allowing external concentrations to vary—important in crowded extracellular spaces where external volume fractions are small—would connect to activity-driven extracellular K^+^ accumulation and spreading depolarization phenomena (85, 86). Finally, while we treated pumps and transporters deterministically, growing evidence indicates that ultraslow mode switching and static disorder can be intrinsic to transport proteins (87–89). A fully stochastic treatment of pumps and exchangers, coupled to voltage and concentration dynamics, may be essential for quantitatively predicting organelle-to-organelle heterogeneity and intermittency.

More broadly, this work emphasizes that “channel noise” in small compartments can be a primary driver of state transitions and homeostatic set points. As optical electrophysiology and organelle-resolved ion sensing mature (14, 34, 37, 38, 90, 91), theories that retain discreteness—single-molecule gating, finite copy numbers, and concentration-driven feedback—will be increasingly necessary for interpretation and for mechanistic inference from data.

## Appendix: Effect of single-channel gating on ion concentrations

For concreteness we consider a persistent Na^+^ current, though the discussion applies to any persistent current. The membrane voltage, *V*_m_, evolves according to:

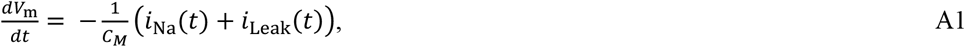

Where *i*_Na_(*t*) is the inward Na^+^ current, *i*_Leak_(*t*) is the outward leak current, and *C*_M_ is the membrane capacitance. The rate of change of concentration of Na^+^ inside the vesicle is:

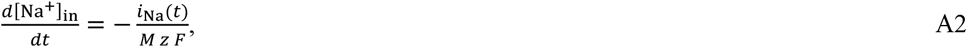

where, *M* is the vesicle volume, *z* is the valence (1 for Na^+^) and *F* is the Faraday constant. The Na^+^ current is:

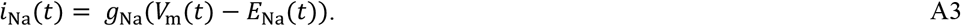

For clarity, we make the time-dependencies explicit and define *g*_x_ as the channel’s open-state conductance and *E*_x_(t) as the channel’s reversal potential; x = Na^+^ or leak ion. We assume that the voltage is in a local equilibrium (i.e. voltage changes are fast compared to the concentrations). Solving Eq. A1 for the steady-state voltage, *V*_ss_(*t*):

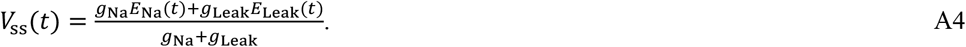

Setting *V*_m_(*t*) = *V*_ss_(*t*) in Eq. A3 and combining Eqs. A2 and A3 gives:

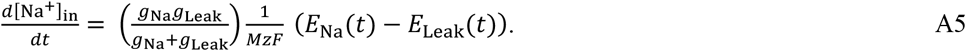

Finally, the sodium reversal potential changes with time as the internal concentration changes:

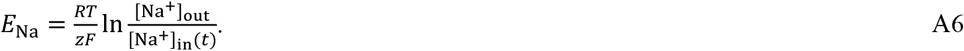

Combining Eqs. A5 and A6 gives an expression for the dynamics of [Na^+^]_in_:

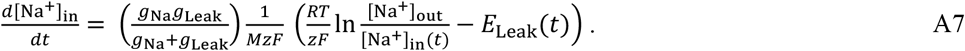

In the case where *g*_Leak_ ∝ *r*^2^ and *M* ∝ *r*^3^ (i.e. a constant leak conductance per unit membrane area in a spherical vesicle), then 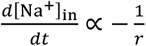 for small vesicles (*g*_Na_≫ *A G*_0_) and 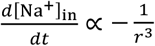 for large vesicles 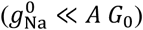.

Equation A7 is nonlinear in [Na^+^]_in_ and does not have a closed-form analytical solution. Numerical integration yields the time-dependent trajectory of [Na^+^]_in_. An approximate timescale can be estimated by extrapolating the initial rate of change to the entire transition from [Na^+^]_in_(0) to [Na^+^]_in_(∞), i.e.

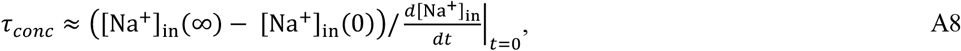

Where

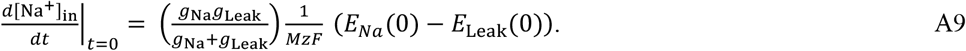

Combining Eqs. A8 and A9 yields Eq. 5 in the main text.

## DATA, MATERIALS, AND SOFTWARE AVAILABILITY

Annotated MATLAB code for stochastic simulations of single-channel vesicles, HH-model vesicles, and macrophage endosomes is provided in a publicly accessible GitHub repository: https://github.com/adamcohenlab/howell2025small-scale-ephys

Simulated data and analysis codes are available upon request.

## AUTHOR CONTRIBUTIONS

M.R.H. and R.J.X. performed the simulations. R.J.X, M.R.H., and A.E.C. wrote the manuscript. A.E.C. envisioned and supervised this project.

## ACKNOWLEDGEMENTS

We thank Prof. Michael Grabe of UCSF for providing us with the Matlab code for the vesicle-type H^+^ ATPase model. This work was supported by a NSF Graduate Research Fellowship (M.R.H.) and NSF Quantum Sensing for Biophysics and Bioengineering (QuBBe) Quantum leap challenge institute (QLCI) grant OMA-2121044. Components of **Fig 1-3, 5** and **Note S2** were created in BioRender. Howell, M. (2025) https://BioRender.com/nm7zk94

