## Supplementary Information for "Electrophysiology in nanoscale compartments"

### Supplemental Information

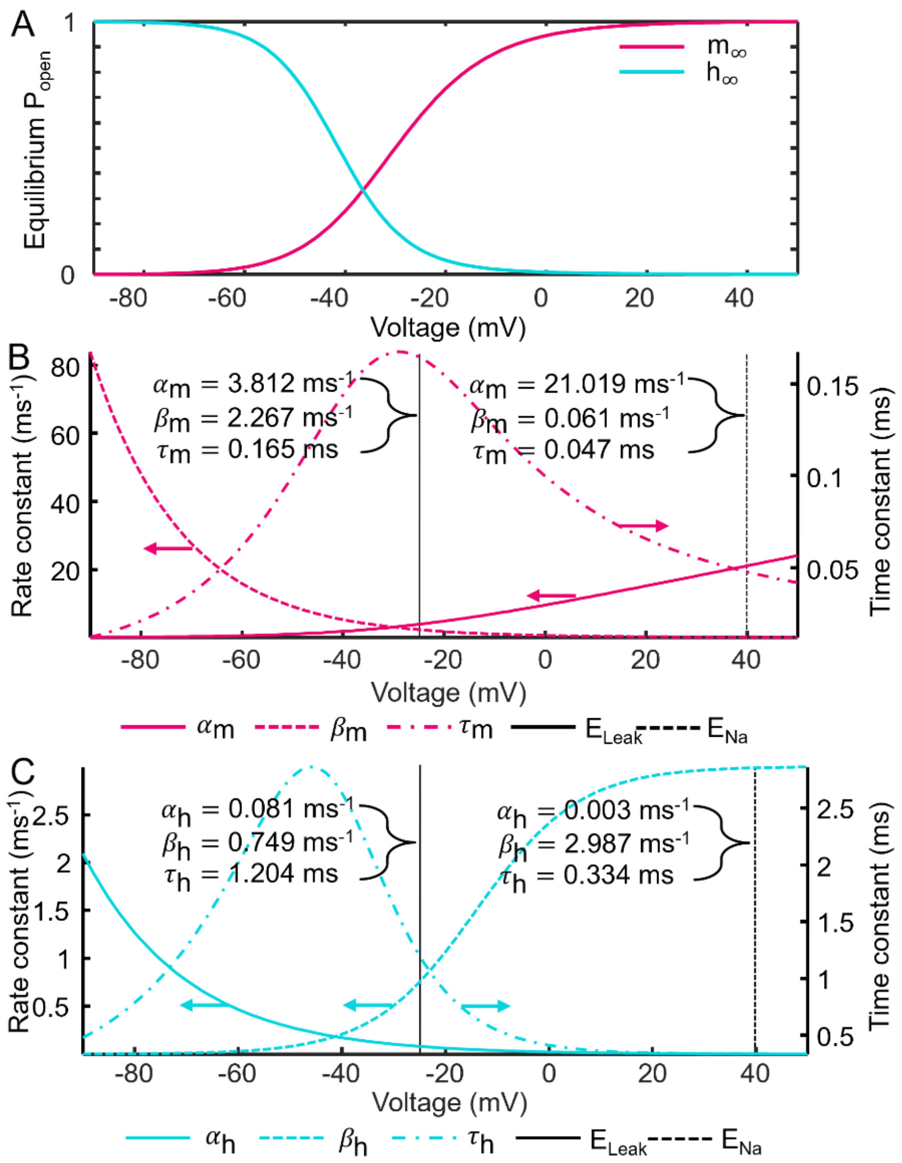


**FIGURE S1. Na$_V$ model.** (**A**) Steady-state activation (pink) and inactivation (cyan). (**B**) Voltage-dependent kinetic parameters of the activation (m) gate. Activation rate, $\alpha_m$ (solid); deactivation rate, $\beta_m$ (dash); time constant $\tau_m$ (dash dot). Vertical lines indicate parameter values at the leak reversal ($E_{Leak}$, solid black line) used in **Fig. 2B-E** and the $Na^+$ reversal potential ($E_{Na}$, dash black line). (**C**) Same as **B**, for the inactivation (h) gate.

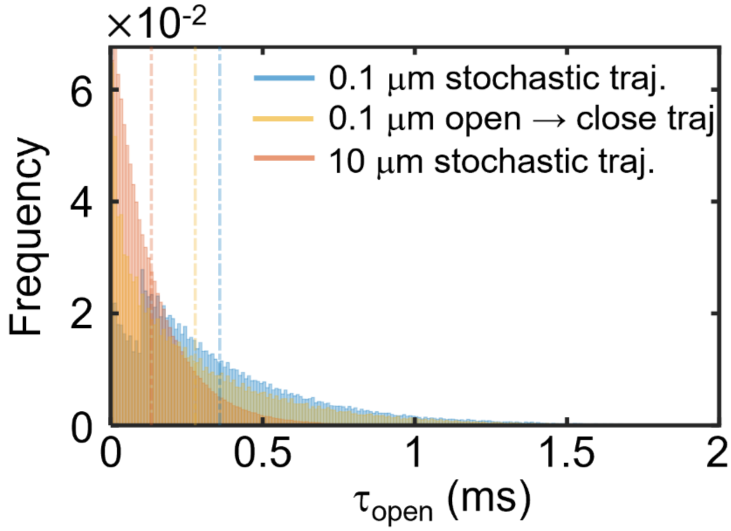


**FIGURE S2. Time-dependent open-state lifetime in single-channel Na$_V$ vesicles.** Open-state dwell time histograms for spontaneous gating trajectories of single Nav channels in vesicles of radius $r = 0.1$ μm (blue) and $r = 10$ μm (orange); simulations 5,000 s. Leak reversal potential, $E_{\text{Leak}} = -25$ mV. Shown for comparison are the open-state lifetimes of Nav in single-channel $r =$ 0.1 μm vesicles were the Nav channel is initialized in the open state at $V_{\text{m}} = E_{\text{Leak}}$ (yellow); $2 \times 10^5$ independent trials. Dashed vertical lines indicate the distribution means; $r = 0.1$ μm stochastic trajectory $\langle \tau_{\text{open}} \rangle = 0.36 \pm 0.33$ ms (blue); $r = 10$ μm stochastic trajectory $\langle \tau_{\text{open}} \rangle =$ $0.13 \pm 0.13$ ms (orange); and $r = 0.1$ μm open $\rightarrow$ closed lifetime $\langle \tau_{\text{open}} \rangle = 0.28 \pm 0.31$ ms (yellow); (mean $\pm$ s.d).

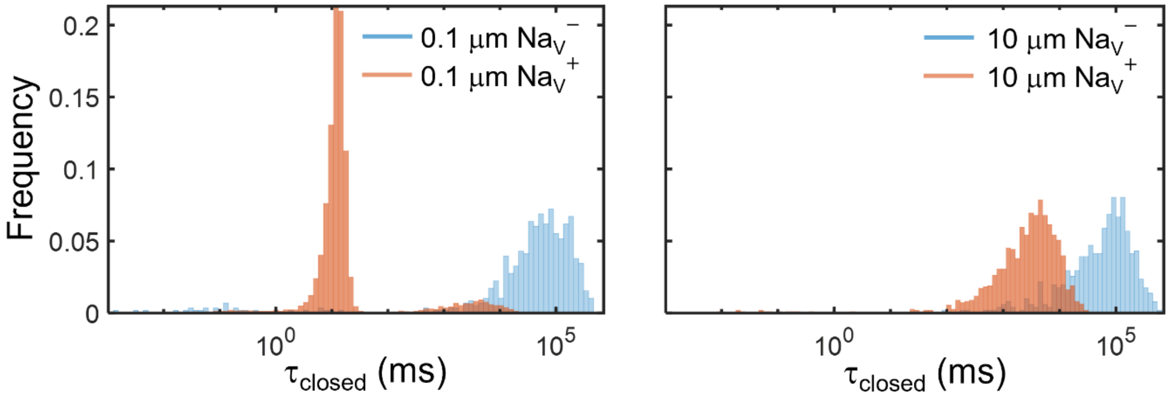


**FIGURE S3. Nanoscale vesicles support distinct patterns of channel gating.** Closed state dwell-time histograms of spontaneous channel gating in single Nav channel vesicles of radii (left) $r = 0.1$ μm and (right) $r = 10$ μm. HH-type Nav gating parameters were adjusted to generate a model with substantial window current (denotated by $Na_V^+$, orange) and a model with negligible window ($Na_V^-$, blue). $Na_V^+$ simulations 5,000 s; $Na_V^-$ results are aggregated from five independent trials of 5,000 s simulations. $E_{leak} = -60$ mV. Modified gating model parameters are described in **Note S1.**

**NOTE S1** Markov model of Nav and Kv gating. Diagram adapted from (1). Abbreviations: I – inactivated; C – closed; O – open.

Nav:

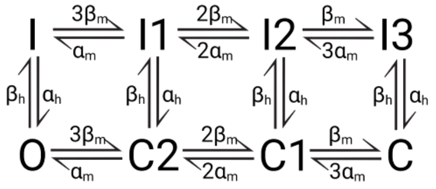


Kv:

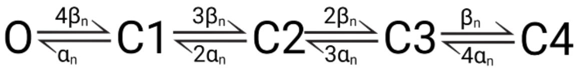


Transition rate functions for default Nav model, adapted from (2), where all times are measured in milliseconds and all voltages in millivolts:

$$\alpha_{\mathrm{m}} = \frac{0.1(V + 30 + S_{\mathrm{m}})}{1 - \exp\left(-\frac{(V + 30 + S_{\mathrm{m}})}{10A_{\mathrm{m}}}\right)}$$

$$\beta_{\mathrm{m}} = 4\exp\left(-\frac{(V + 55 + S_{\mathrm{m}})}{18A_{\mathrm{m}}}\right)$$

$$\alpha_{\mathrm{h}} = 0.07\exp\left(-\frac{(V + 44 + S_{\mathrm{h}})}{20A_{\mathrm{h}}}\right)$$

$$\beta_{\mathrm{h}} = \frac{1}{1 + \exp\left(-\frac{(V + 14 + S_{\mathrm{h}})}{10A_{\mathrm{h}}}\right)}$$

$$\alpha_{\mathrm{n}} = \frac{0.01(V + 34)}{1 - \exp\left(-\frac{(V + 34)}{10}\right)}$$

$$\beta_{\mathrm{n}} = 0.125\exp\left(-\frac{(V + 44)}{80}\right)$$

Default Nav contains moderate window currents: $S_m = S_h = 0$; $A_m = A_h = 1$.

Nav with large window currents ($Nav^+$): $S_m = 10$; $S_h = -10$; $A_m = 0.55$; $A_h = 0.55$.

Nav with small window currents ($Nav^-$): $S_m = -10$; $S_h = 19$; $A_m = 0.95$; $A_h = 1$.

**NOTE S2** Nav1.5 state diagram and transition rate functions from (3). The unified model developed by Balbi *et al.* accounts for Nav1.1-1.9 dynamics using fewer channel states than the Markov model of HH-type Nav (**Note S1**) and isoform-specific gating rates.

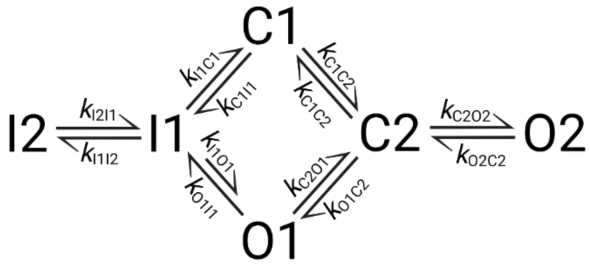


Abbreviations: I – inactivated; C – closed; O – open. All times are measured in milliseconds and all voltages in millivolts:

$$k_{\mathrm{C1C2}} = \frac{10}{1 + e^{\frac{V-(-13)}{-10}}}$$

$$k_{\mathrm{C2C1}} = \frac{1}{1 + e^{\frac{V-(-43)}{8}}} + \frac{10}{1 + e^{\frac{V-(-13)}{-10}}}$$

$$k_{\mathrm{C2O1}} = \frac{10}{1 + e^{\frac{V-(-23)}{-10}}}$$

$$k_{\mathrm{O1C2}} = \frac{1}{1 + e^{\frac{V-(-53)}{8}}} + \frac{10}{1 + e^{\frac{V-(-23)}{-10}}}$$

$$k_{\mathrm{C2O2}} = \frac{0.05}{1 + e^{\frac{V-(-10)}{-10}}}$$

$$k_{\mathrm{O2C2}} = \frac{2}{1 + e^{\frac{V-(-50)}{10}}} + \frac{0.08}{1 + e^{\frac{V-(-20)}{-10}}}$$

$$k_{\mathrm{O1I1}} = \frac{7}{1 + e^{\frac{V-(-44)}{13}}} + \frac{10}{1 + e^{\frac{V-(-19)}{-13}}}$$

$$k_{\mathrm{I1O1}} = \frac{1 \times 10^{-5}}{1 + e^{\frac{V-(-20)}{10}}}$$

$$k_{\mathrm{I1C1}} = \frac{0.19}{1 + e^{\frac{V-(-110)}{7}}}$$

$$k_{\mathrm{C1I1}} = \frac{0.016}{1 + e^{\frac{V-(-92)}{-6}}}$$

$$k_{\mathrm{I1I2}} = \frac{2.2 \times 10^{-4}}{1 + e^{\frac{V-(-50)}{-5}}}$$

$$k_{\mathrm{I2I1}} = \frac{1.8 \times 10^{-3}}{1 + e^{\frac{V-(-90)}{30}}}$$

**TABLE S1** Electrophysiological parameters and initial conditions of the HH neuron-type vesicle model.

| Parameter | Value | Reference |
|---|---|---|
| Radius (μm) | 0.02 - 10 | |
| Nav channel density ($\mu m^{-2}$) | 0.8 - 8000 | |
| Kv channel density ($\mu m^{-2}$) | 0.2 - 2000 | |
| Extravesicular volume ($\mu m^{-3}$) | 100000 | |
| $C_m$, Membrane capacitance ($pF/\mu m^{-2}$) | 0.01 | (2) |
| $g_{Na_V}$ , Nav single-channel conductance (nS) | 0.014 | (4) |
| $g_{K_V}$ , Kv single-channel conductance (nS) | 0.020 | (5) |
| $g_{Na^+,leak}$ , $Na^+$ leakage conductance ($nS/\mu m^{-2}$) | 0.000175 | (2) |
| $g_{K^+,leak}$ , $K^+$ leakage conductance ($nS/\mu m^{-2}$) | 0.0005 | (2) |
| $g_{Cl^-,leak}$ , $Cl^-$ leakage conductance ($nS/\mu m^{-2}$) | 0.0005 | (2) |
| $I_{pump,max}$ , Maximum v-ATPase pump rate (pA) | 0.0525 | (2, 6) |
| $[Na^+]_i$ , Intravesicular $Na^+$ concentration (mM) | 27 | (2) |
| $[Na^+]_e$ , Extravesicular $Na^+$ concentration (mM) | 120 | (2) |
| $[K^+]_i$ , Intravesicular $K^+$ concentration (mM) | 131 | (2) |
| $[K^+]_e$ , Extravesicular $K^+$ concentration (mM) | 4 | (2) |
| $[Cl^-]_i$ , Intravesicular $Cl^-$ concentration (mM) | 9.66 | (2) |
| $[Cl^-]_e$ , Extravesicular $Cl^-$ concentration (mM) | 124 | (2) |
| $V_m$ , Initial membrane voltage (mV) | -68.0 | |
| $E_{Na^+}$, $Na^+$ reversal potential (mV) | 39.7 | |
| $E_{K^+}$, $K^+$ reversal potential (mV) | -92.9 | |
| $E_{Cl^-}$ , $Cl^-$ reversal potential (mV) | -68.0 | |

**TABLE S2** Electrophysiological parameters and initial conditions of the endosome model.

| Parameter | Value | Reference |
|---|---|---|
| Radius (μm) | 0.2 | |
| Nav1.5 channel density ($\mu m^{-2}$) | 0 - 6400 | |
| v-ATPase pump density ($\mu m^{-2}$) | 700 | (7) |
| ClC-7 channel density ($\mu m^{-2}$) | 300 | (8) |
| Extravesicular volume ($\mu m^{-3}$) | 100000 | |
| $C_m$, Membrane capacitance ($pF/\mu m^{-2}$) | 0.01 | (2) |
| $g_{Nav\ 1.5}$, Nav1.5 single-channel conductance (nS) | 0.0173 | (9) |
| $[Na^+]_i$ , Luminal $Na^+$ concentration (mM) | 100 | (10) |
| $[Na^+]_e$ , Intracellular $Na^+$ concentration (mM) | 12 | |
| $[Cl^-]_i$ , Luminal $Cl^-$ concentration (mM) | 19 | (10) |
| $[Cl^-]_e$ , Intracellular $Cl^-$ concentration (mM) | 10 | |
| $pH_i$ , Luminal pH | 6.2 | (10) |
| $pH_e$ , Intracellular pH in macrophages | 7.5 | (11) |
| Luminal $H^+$ buffering capacity (mM/dpH) | 40 | (7) |
| $V_m$ , Initial membrane voltage (mV) | 19 | (12) |
| $E_{Na^+}$, $Na^+$ reversal potential (mV) | -56.5 | |
